# CRISPR-mediated knock-in of transgenes into the malaria vector *Anopheles funestus*

**DOI:** 10.1101/2021.03.31.437891

**Authors:** Charlotte Quinn, Amalia Anthousi, Charles Wondji, Tony Nolan

## Abstract

The ability to introduce mutations, or transgenes, of choice to precise genomic locations has revolutionised our ability to understand how genes and organisms work.

In many mosquito species that are vectors of various human disease, the advent of CRISPR genome editing tools has shed light on basic aspects of their biology that are relevant to their efficiency as disease vectors. This allows a better understanding of how current control tools work and opens up the possibility of novel genetic control approaches, such as gene drives, that deliberately introduce genetic traits into populations. Yet for the *Anopheles funestus* mosquito, a significant vector of malaria in sub-Saharan Africa and indeed the dominant vector species in many countries, transgenesis has yet to be achieved.

We describe herein an optimised transformation system based on the germline delivery of CRISPR components that allows efficient cleavage of a previously validated genomic site and preferential repair of these cut sites via homology-directed repair (HDR), which allows introduction of exogenous template sequence, rather than end-joining repair. The rates of transformation achieved are sufficiently high that it should be able to introduce alleles of choice to a target locus, and recover these, without the need to include additional dominant marker genes. Moreover, the high rates of HDR observed suggest that gene drives, which employ an HDR-type mechanism to ensure their proliferation in the genome, may be well suited to work in *An. funestus*.

## INTRODUCTION

Mosquitoes belonging to the genus *Anopheles* are vectors of a range of agents of human disease, including viruses, parasitic worms and, most notably, malaria parasites. A way to introduce defined genetic changes precisely into these insects can drastically improve our understanding of the genes responsible for key traits that determine its capacity as a disease vector, such as parasite susceptibility, female reproductive output, human biting behaviour, as well as helping in elucidating mechanisms of resistance to insecticides. Moreover, the ability to introduce genes of choice opens up the possibility of genetic control – the deliberate introduction of traits into a mosquito population to reduce its capacity to transmit disease.

To date, most attention has focused on two members of the *Anopheles gambiae* species complex, *Anopheles gambiae (s*.*s*.) and *Anopheles coluzzi*, that are the dominant vectors of human malaria across most of sub-Saharan Africa, where malaria burden is highest. However, *Anopheles funestus*, not a member of this species complex, is the dominant vector in many places, notably in southerly regions of sub-Saharan Africa, and insecticide resistance in this species has been associated with local malaria resurgence there and loss of efficacy of control interventions (Cohuet et al. 2004; Riveron et al. 2019). Since the first demonstrations of a working, transposon-based transgenic technology for mosquitoes over 20 year ago, most of the major mosquito vectors have now been transformed (Grossman et al. 2001; Catteruccia et al. 2000; Coates et al. 1998; Jasinskiene et al. 1998; Allen et al. 2001), yet *An. funestus* still remains the exception. This may in part simply reflect the logistical difficulty associated with rearing this species in the laboratory (Hunt et al. 2005; Ngowo et al. 2021; Morgan et al. 2010), but may also reflect a quirk in its biology that make it more refractory to the process of germline transformation.

The prospects for transforming a mosquito species of choice have been greatly improved by the general applicability of CRISPR-Cas9 genome engineering tools to a wide range of species (Hammond et al. 2016; Gantz et al. 2015; Purusothaman et al. 2021). CRISPR-mediated cleavage can be repaired via one of two pathways: non-homologous end-joining (NHEJ) repair and homology-directed repair (HDR). Recently, CRISPR-mediated cleavage was used to introduce random, heritable mutations via NHEJ at specific sites in the genome of *An. funestus* (Li, Akbari, and White 2018). However, to date there still has been no documented report of HDR introduction (often referred to as ‘knock-in’) of either transgenes, or specified pre-determined mutations, into this species. This method relies on the provision of a DNA ‘donor’ template containing regions of homology either side of the cut site, designed in such a way that the cell’s HDR machinery uses it as a template for repair, resulting in the incorporation of an allele or transgene of choice. A method of this type is crucial for two reasons: it would allow confirmation of genetic factors/signatures identified in field populations that putatively confer important phenotypes, such as insecticide resistance, with strong relevance to control programmes; it opens the door to novel genetic control approaches, such as gene drive, which result in the specific introduction and spread of transgenic traits into mosquito populations as a form of vector control.

The relative propensity for CRISPR-induced DNA breaks to be repaired by HDR or NHEJ can show strong variation according to whether the tissue is of somatic or germline origin, and even within the germline it can be dependent on the level and timing of Cas9 expression (Kandul et al. 2020; Li et al. 2017; Hammond et al. 2021). For the purposes of generating stable transformed mosquito strains it is essential that the CRISPR-induced modifications occur in the germline, so that they are transmitted to their offspring from which a modified line can be established. We have previously demonstrated high rates of CRISPR-mediated HDR in generating transgenic *An. gambiae* by microinjecting embryos with a plasmid-based source of Cas9 under a germline-specific promoter together with a donor template designed to incorporate a dominant marker gene. Given this success in *An. gambiae*, we investigated whether a similar approach could work in *Anopheles funestus*, using components likely to work broadly across both species.

Our experimental design allowed us to determine the relative rates of NHEJ compared to HDR following Cas9 activity in injected embryos. We were able to show rates of HDR-mediated germline transformation comparable to those achieved for *An. gambiae*, suggesting that the wealth of functional genetics tools and options for genetic control that exist in that species may soon be available in *Anopheles funestus*.

## RESULTS

As a proof of principle, we determined to target the *An. funestus* X-linked gene *white*, which when mutated produces a visible phenotype in the eye (Li et al. 2017), with an HDR template designed to introduce a dominant fluorescent marker gene expressing cyan fluorescent protein (eCFP) (Fig. 1a). We produced a ‘helper’ plasmid containing the Cas9 gene under the control of the germline-specific promoter of the *An. gambiae vasa* gene (Papathanos et al. 2009). Given that this promoter sequence from *An. gambiae* manages to confer germline expression even in the distantly related mosquito *Aedes aegypti* (Akbari et al. 2014), we reasoned that it should work similarly in *An. funestus*. The helper plasmid also contained a construct producing a *white*-specific guide RNA under transcriptional control of the U6 promoter (Hammond et al. 2016).

**Figure 1.**
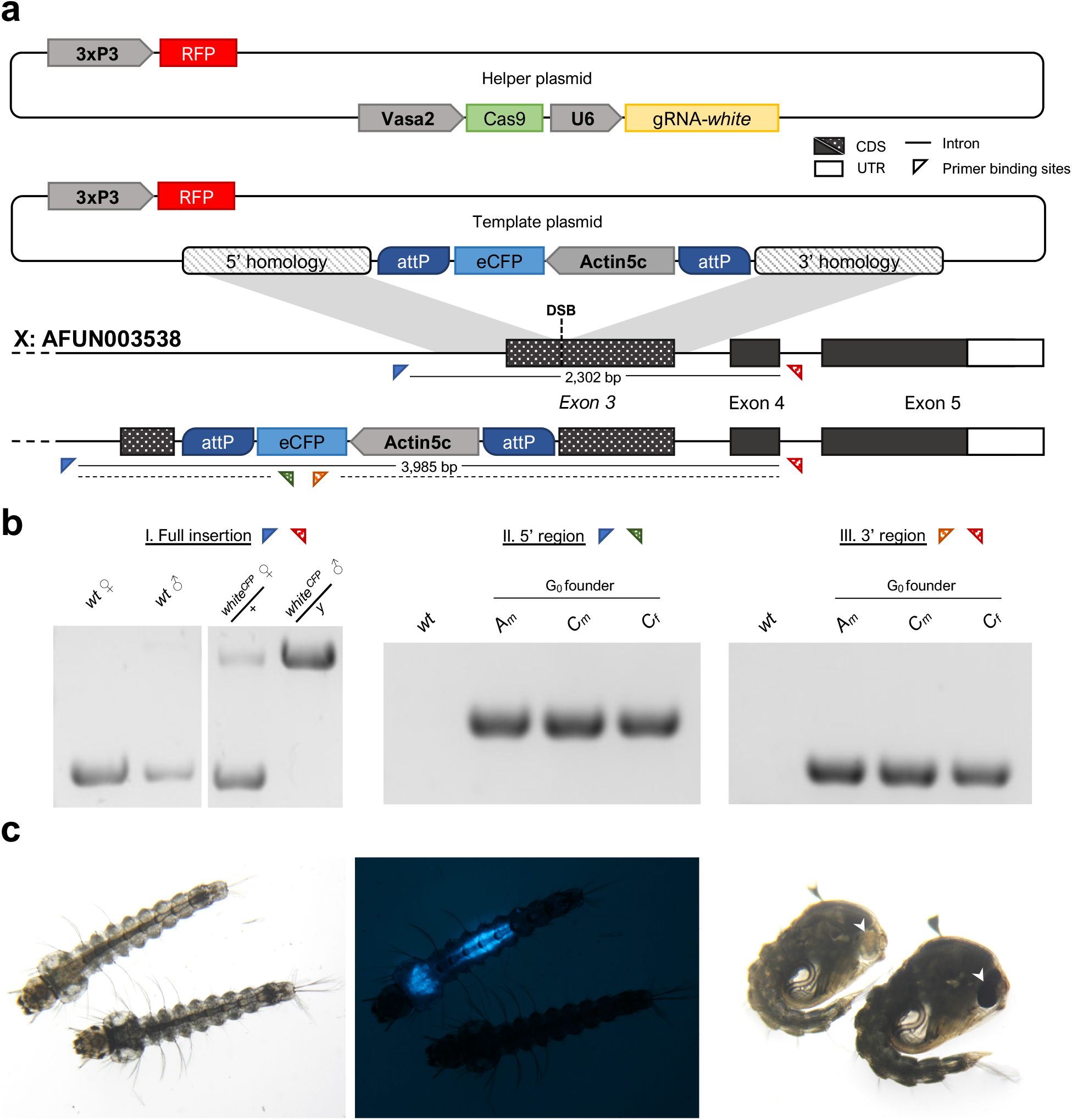
Targeted disruption of the *white* gene (AFUN003538) by CRISPR-mediated homology-directed repair. (**a**) Schematic representation of the HDR knock-in process. DNA repair is mediated through the concurrent microinjection of two plasmid assemblies: a ‘helper’ plasmid, designed to induce a double-stranded break (DSB) at the target locus upon expression, containing a source of Cas9 and gRNA under the control of the vasa2 and U6 promoters respectively; and a ‘template’ plasmid containing the insert region (an actin5c::eCFP cassette enclosed within two reversible ϕC31 *attP* recombination sequences) flanked both 5’ and 3’ by regions of homology approximately 1kbp upstream and downstream of the cut site. (**b**) Diagnostic PCRs of transgenic offspring (G2) deriving from different founders. PCR primer binding sites are represented by triangles). ‘External’ primers flanking the full insertion and binding outside the regions of homology included in the donor construct (red and blue triangles, I) are used to discriminate between wild-type individuals and females heterozygous or males hemizygous for the *white*^*CFP*^ allele. ‘Internal’ primers complementary to the knocked-in eCFP sequence (green and orange triangles) are used with external primers to amplify the 5’ region upstream (II) or 3’ region downstream (III) of the eCFP cassette. Three separate transgenic lines were produced and assigned according to injection set (A - D) and sex of founder individual or group (m – males or f – females). wt – wild-type control. (**c**) Bright-field (left and right) or standard fluorescent (middle) microphotographs of representative individuals demonstrating either the wild-type or *white*^*CFP*^ mutant phenotype. Under bright-field illumination, mutant larvae (left) and pupae (right) exhibit lighter global pigmentation compared to wild-type, and are white-eyed (arrows). Transgenic *white*^*CFP*^ individuals additionally express cyan fluorescent protein from the actin5C promoter, in a pattern (lower midgut and gastric caecae) consistent with the activity of this promoter in other Anopheles species (middle).

Together with the helper plasmid we included an HDR ‘donor’ construct (Fig. 1a) that contained, as a dominant marker the gene encoding an enhanced version of the cyan fluorescent protein (eCFP) under the transcriptional control of the actin5c promoter from *Drosophila melanogaster* which has previously been shown to work well in *An. stephensi* and *An. gambiae* (Catteruccia et al. 2000; Bernardini et al. 2014). Again, given this demonstration of conservation of the promoter activity across such a large evolutionary distance, coupled with the ubiquitous use of eCFP as a visual reporter in a range of species, we reasoned that these components would function as a dominant marker cassette in *An. funestus*.

We injected *An. funestus* eggs of the FANG strain (Hunt et al. 2005) following a standard microinjection protocol (Fuchs, Nolan, and Crisanti 2012; Benedict 2007) containing only minor modifications. Over four separate sets of injections we aligned approximately 1700 embryos for injection. Among the embryos surviving the procedure we screened for those larvae showing visual expression of either the CFP or RFP marker genes transiently from either the helper or donor plasmids. Encouragingly, we were able to see either RFP or CFP expression in 44 of 223 surviving larvae (∼19%, Table 1), indicating that these marker genes, and their respective transcriptional units, are suitably expressed in *An. funestus* cells, at least from an episomal source of injected plasmid. Since this fraction of larvae (which we term ‘transient’ G_0_) are sure to have been injected successfully with a considerable quantity of the transformation plasmids, and given that *An. funestus* can be cumbersome to rear and cross generally, we focused the majority of our efforts on maximising the screening of their progeny, rather than the ‘non-transients’ that may represent non-injected, or sub-optimally injected, G_0_ individuals.

**Table 1.** Outcome of independent injection experiments (sets A – D).

| Injection Set | Embryos aligned | Larvae surviving | G<sub>0</sub> Larvae with transient CFP or RFP expression | G<sub>0</sub> transient individuals surviving to adulthood | G<sub>0</sub> adults with eye colour mosaicism | Crossing scheme | Egg batches produced | HDR events/total G1 (%) |
| --- | --- | --- | --- | --- | --- | --- | --- | --- |
| A | 640 | 53 | 11 | 4 | 0 | Transient ♂(3) x wt outcross | 5 | 71/174 (42.5) |
|  |  |  |  |  |  | Transient ♀(1) wt outcross | 0 | 0/0 (0) |
|  |  |  |  |  |  | NT♂/ ♀ intercross | 1 | 0/130 (0) |
| B | 350 | 68 | 7 | 4 | 0 | Transient ♂(2) x wt outcross | 1 | 0/21 (0) |
|  |  |  |  |  |  | Transient ♀(2) x wt outcross | 0 | 0/0 (0) |
|  |  |  |  |  |  | NT♂/ ♀ intercross | 1 | 0/55 (0) |
| C | 420 | 40 | 12 | 7 | 0 | Transient ♂(5) x wt outcross | 4 | 90/564 (16.0) |
|  |  |  |  |  |  | Transient ♀(2) x wt outcross | 4 | 20/140 (14.3) |
|  |  |  |  |  |  | NT♂/ ♀ intercross | 1 | 0/61 (0) |
| D | 300 | 72 | 14 | 11 | 0 | Transient ♂ (5) x wt outcross | 0 | 0/0 (0) |
|  |  |  |  |  |  | Transient ♀(6) x wt outcross | 2 | 0/73 (0) |
|  |  |  |  |  |  | NT♂/ ♀ intercross | 1 | 0/75 (0) |

We crossed G_0_ individuals to wild type and screened for the presence of CFP-positive offspring – suggestive of a HDR-targeting event in the germline of the parent. Across the injection sets, from a total of 26 surviving adult G_0_ individuals, we recovered 3 separate founders that produced CFP-positive offspring (Fig 1c), representing a transformation rate of ∼11%, considering this class only (or ∼3% considering all G_0_, transient and non-transient). Among the offspring of these founders the ratio of transgenic to non-transgenic offspring was relatively high, ranging from 14% to 43% (Table 1), suggesting that the transformation event happens early in germline development when there are relatively few germline stem cells. The fact that most individuals were CFP-positive only (and not RFP-positive also) is consistent with CRISPR-mediated HDR integration of the CFP cassette contained within the regions of homology upstream and downstream of the target site. To confirm this, we performed a diagnostic PCR using primers that bind the genomic target region externally to the regions of homology contained in the donor plasmid (Fig 1a). The presence of the complete CFP cassette at the target locus and the precise nature of the integration at the 5’ and 3’ end of the construct was confirmed (Fig 1b), and we named the allele thus generated as *white*^*CFP*^. Our expectation was that the *white*^*CFP*^ allele would represent a null allele, given that indels at this target site result in a white-eye phenotype (Li, Akbari, and White 2018). Indeed, since the *white* gene resides on the X chromosome, hemizygous *white*^*CFP*^ males had white eyes whereas heterozygous females exhibited a wild type eye colour, confirming that the *white*^*CFP*^ allele is both recessive and, likely, null. Among the G_1_ progeny we also recovered a small fraction of offspring (3.8% of all transgenic offspring) that were both RFP+ and CFP+, suggesting that in rare cases semi-legitimate recombination may occur, inserting at least a section of the donor plasmid external to the homology-flanked cassette that includes an RFP cassette in the plasmid backbone (Fig 1a). However, we did not investigate these rare events molecularly. The *white*^*CFP*^ HDR allele showed normal Mendelian inheritance and, in homozygosity in females, resulted in a white eye phenotype (Table S1). One of our motivations for using a plasmid-based source of Cas9 under a germline-specific promoter was to target Cas9 activity to the germline where rates of HDR seem to be very high, in *Anopheles* species at least, and to have a system that could minimise somatic activity of the Cas9. Previously, it was shown that while injection of a mix of Cas9 protein and gRNA could generate indels at the target site in the germline, it also generated a mosaic of somatic mutations in these individuals, including the mutation of both alleles (bi-allelic mutations) in some cells, resulting in eye colour mosaicism in these G_0_ individuals (Li, Akbari, and White 2018). On the contrary, in the G_0_ individuals injected with the vasa-driven source of Cas9 we failed to observe any eye colour mosaicism in males or females (Table 1). A T7 assay revealed that rates of end-joining mutations were very low in injected individuals, to the point where they were barely detectable (Supp Fig 1). Indeed, in one founder that produced HDR-mediated transgenic offspring we failed to observe any signal of end-joining activity in the T7 assay. Moreover, we also found no evidence of any germline transmission of Cas9 generated indels from any of the founders – these indels would be expected to result in a non-CFP, white-eyed phenotype in hemizygous males, yet we failed to observe any such events among the siblings of HDR-mediated transgenic offspring. Taken together, these data support the hypothesis that this transformation protocol, using a germline-restricted source of Cas9, strongly favours the generation of HDR-mediated repair over end-joining repair.

## DISCUSSION

This is the first report of the generation of transgenic *Anopheles funestus*. The availability of a CRISPR-based technology to introduce transgenes, or to introduce precise and defined changes, at specific genomic loci, has the potential transform our ability to perform functional genetics in insects, such as exploring the function of genes putatively controlling key phenotypes such as insecticide resistance (Samantsidis et al. 2020; Itokawa et al. 2016), investigating genes that are essential for important insect behaviours (Raji et al. 2019) or identifying genes that are important for key traits such as immunity and female reproductive capacity (Hammond et al. 2016).

For *An. funestus*, a pressing aspect to investigate is the nature of its resistance to pyrethroid insecticides, a mainstay of insecticide-treated bednets, where the resistance mechanism appears to be different to that observed in *An. gambiae* and may involve copy number variation, allelic variation of coding regions and/or the insertion of cis-regulatory sequences that lead to upregulation of expression (Mugenzi et al. 2019; Weedall et al. 2019; Weedall et al. 2020; Ibrahim et al. 2015). A CRISPR-based HDR tool will be crucial in investigating these hypotheses.

The high frequency with which transgenics were recovered using our approach mean that it should be possible, by judicious molecular genotyping of offspring, to recover precisely modified mosquitoes without the need for insertion of a dominant marker gene, that might have confounding effects on phenotype. Of course, the efficiency of targeting may vary according to gRNA:locus combination, and additional future experiments it will be important to confirm targeting rates across a range of loci. The relative rates of HDR vs end-joining repair following Cas9-induced DNA breaks can vary according to cell type and between germline and soma (Lin et al. 2014; Hammond et al. 2021; Kandul et al. 2020). Additionally, there appears to be variation in the rates of CRISPR-based transgenesis between species with the rates for *Aedes aegypti*, for example, generally being significantly lower than those observed in *Anopheles gambiae* (Kistler, Vosshall, and Matthews 2015; Hammond et al. 2016). However, it is difficult to disentangle effects due to differing biology between the species and effects due to different transformation systems/setups. Indeed, the HDR transformation rates were improved several fold in *Aedes aegypti* by injecting a line containing a transgenic source of germline Cas9 (Li et al. 2017). Our approach, of injecting a plasmid-based source of Cas9 active in the germline coupled with a gRNA transcription unit has several attractive features: it obviates the need to inject a Cas9-expressing transgenic line; the plasmid-based DNA solution is much less labile than protein/RNA-based mixtures; rates of HDR are high; and the relative ratio of HDR:NHEJ events is very high, as evidenced by the almost complete lack of somatic mosaicism observed after injection. This latter aspect is particularly important if the target gene is essential and extensive mosaicism might reduce the recovery of desired events.

Finally, an important consideration for the plausibility of generating gene drives, which are designed to bias their inheritance among offspring and thereby spread linked traits into a population (Alphey et al. 2020), are the relative rates of HDR vs NHEJ. Homing-based gene drives (Burt 2003; Esvelt et al. 2014) have, to date, shown the most promise in terms of development and proof-of-principle demonstration in the laboratory, in a number of mosquito species (Li et al. 2020; Hammond et al. 2016; Gantz et al. 2015). This type of gene drive encodes a Cas9:gRNA unit that cleaves germline chromosomes at an endogenous target sequence and hijacks the HDR pathway to copy itself into the repaired site, thereby increasing its copy number. Thus, HDR is essential for this type of gene drive to work. On the flip side, NHEJ repair events following gene drive activity not only reduce the probability of the drive copying itself directly but the small indels can generate target sites that are resistant to future gene drive targeting (Champer et al. 2017; Hammond et al. 2017). Therefore, the high rates of HDR that we observed in *An. funestus* augur well for the prospect of developing gene drives for control of this considerable vector of malaria that, coupled now with a capacity to perform functional genetics, should expand the range of innovative control tools available and augment those that already exist.

## MATERIALS AND METHODS

### Generation of transformation constructs for gene targeting by CRISPR

The helper plasmid, containing source of Cas9 and guide RNA, was a derivative of plasmid p165 (accession ID: <u>KU189142</u>) (Hammond et al. 2016), containing a vasa2::SpCas9 construct and a U6::gRNA cassette containing a spacer cloning site based on Hwang et al. (2013). Oligos (forward – TGCTGGTGAGCTCCTTGCGGTGAT; reverse – AAACATCACCGCAAGGAGCTCACC) were designed to make a functional spacer targeting a previously validated target site in the *Anopheles funestus white* gene (AFUN003538) (Li, Akbari, and White 2018) were annealed and cloned via *Bbs*I Golden Gate cloning into the spacer cloning site to produce helper plasmid p18.

The HDR ‘donor’ construct contained regions of homology approximately 1 kb long immediately upstream and downstream of the target cut site, amplified from pooled genomic DNA of *Anopheles funestus* mosquitoes (FANG strain, LSTM) with primer sets designed for Gibson assembly of the two homology arms flanking a dominant marker cassette, designed to express eCFP. Primer sequences (sequences for Gibson cloning in small case) for the homology arms were as follows: 5’ homology, forward – gcgagctcgaattaaccattgtggAACCGTGCCTCTATTCTCAGC; 5’ homology, reverse – tggggtaccggtACCGCAAGGAGCTCACCAC; 3’ homology, forward – atcctgaacgcGTGATGGGTAGTTCCGGTGC; 3’ homology, reverse – tactccacctcacCCATGGGACCCAACATCGGATCCTTCAGAAC. The dominant marker cassette, containing an enhanced cyan fluorescent protein (eCFP) unit under the promotional control of the *Drosophila Actin5c* promoter (Han, Levine, and Manley 1989) enclosed within two □C31 attP recombination sequences, was amplified from plasmid pK104c (gift of K. Kyrou) using primers kk047 (5’-ACCGGTACCCCAATCGTTCA-3’) and kk048 (5’-ACGCGTTCAGGATTATATCT-3’). The homology arms and marker cassette were cloned by Gibson assembly into *Mlu*I- and *Msh*TI-digested plasmid K103 (K. Kyrou), to generate the final donor plasmid (p19).

### *An. funestus* rearing – Insectary Conditions, Microinjections, Screening of Transgenics

A large (>5,000 adults) colony of *An. funestus* mosquitoes, FANG strain (Hunt et al. 2005) is routinely held in insectaries at the Liverpool Insect Testing Establishment (LITE), a dedicated GLP-approved facility, adjoined to our host institution, for the continuous production of mosquitoes and the testing of vector control products. The FANG strain originated from Angola and is fully susceptible to all insecticides (Hunt et al. 2005). For general maintenance, FANG colonies at LITE were provided a blood meal twice prior to oviposition, using a Hemotek Membrane Feeding System (Hemotek Ltd., Blackburn, UK), and with screened human blood procured from NHS bloodbanks. In brief, eggs were floated in purified water with 0.01% w/v yeast hatching solution and reared at larval densities of 0.24 larvae/mL. Larvae were fed increasing amounts of ground TetraMin® Tropical Flakes under the following regime: day 1 (day of hatching) ∼ 100 µg/larvae; then day 2–7 ∼ 200 µg/larvae; then day 8–10 ∼ 233 µg/larvae, with day 10 being the first day of pupation.

Prior to microinjection a cohort of mosquitoes (approx. 1500 adults) were maintained in dedicated insectary facilities adjacent to the microinjection facility, at a temperature of 26 ± 2 °C, a relative humidity of 70 ± 10%, and under an 11-hour light/dark cycle with 1-h dawn/dusk transitions. Female mosquitoes were provided a blood meal from a human volunteer in accordance with LSTM’s code of practice for arm-feeding *Anopheles* mosquitoes. Gravid females were encouraged to oviposit 60–96h hours after blood feeding by introducing them into oviposition ‘chambers’, comprising of a 50ml falcon tube with the conical base removed, covered at one end with dual-layered netting to create a secure way of entry, and plugged at the other end with a shallow, concave plastic cap. Mosquitoes were introduced inside the chambers in small groups (of approximately 10-15 individuals) and left to acclimatise in the dark for at least 30 minutes before initiating oviposition by adding 2ml distilled H_2_O to the cap.

Embryos were injected using a modified version of the method of Benedict (2007) as described previously (Fuchs, Nolan, and Crisanti 2012). Briefly, embryos were aligned against semi-moist nitrocellulose paper, wet with distilled water, and injected into the posterior pole at an oblique angle. The injection mix containing helper (100ng/ul) and donor plasmids (300ng/ul) was resuspended in 1x injection buffer (0.1mM sodium phosphate buffer pH6.8; 10mM KCl). Following injection, eggs were floated in 1L distilled water and 0.01% w/v yeast solution, in plastic trays line with filter paper around the waterline to prevent embryos adhering to the plastic above the waterline and dessicating. Surviving G_0_ progeny were screened for transient expression of eCFP or RFP at the 1st larval instar and reared separately, accordingly as ‘transients’ or ‘non-transients’. We found the injection of *An. funestus* embryos slightly more challenging in terms of needle entry and survival, compared to *An. gambiae* and *An. stephensi*. Since it is certain that transient individuals were injected with a significant amount of injection mix, we prioritised our efforts to maximise the screening of the progeny of these individuals as opposed to the non-transient larvae that may include a significant amount of non-injected, or sub-optimally injected, individuals.

Early-stage larvae of the 1^st^ and 2^nd^ instars were fed 0.015% TetraMin® Baby fish food daily; 3^rd^ and 4^th^ instar larvae were provided with 0.02% ground TetraMin® Tropical Flakes. Once transgenic colonies were established, larval trays were maintained at a density of ≤100-150 larvae per tray of size 30cmx30cm.

G_0_ mosquitoes exhibiting transient fluorescent expression as 1^st^ instar larvae were reared in separate groups of males or females and mated to wild-type for ≥7 days before blood-feeding. Subsequent generations were screened for *actin5c::eCFP* expression as mid-stage larvae; *white-*eye phenotypes and sex were assessed at pupal stages. For G_1_, all *white*^*CFP*^ offspring were female and thus mated to wild-type males for ≥7 days before blood-feeding. For G_2_, *white*^*CFP*^ offspring were outcrossed to wild-type in separate groups of males or females. For G_3_, *white*^*CFP*^ offspring from *white*^*CFP*^ G2 males (all female) were outcrossed to wild-type, while *white*^*CFP*^ offspring from *white*^*CFP*^ G2 females (males and females) were intercrossed together. Finally at G_4_, homozygous (white-eye, *white*^*CFP*^ females) were crossed to *white*^*CFP*^ males in order to maintain transgene purity in following generations.

### T7 Endonuclease Assay

To assess for non-HDR mutagenic activity, G_0_ mosquitoes, exhibiting transient fluorescent expression of the helper and donor plasmids, were first assessed for white-eye phenotypes under standard brightfield microscopy. The genomic DNA (gDNA) of individual transient G_0_ mosquitoes was extracted using the Wizard® Genomic DNA Purification Kit (Promega). In the T7 Endonuclease I (T7EI) assay, a 744 bp region spanning the cut-site was first amplified by PCR (primer sequence forward – GGCTGGTGTATGGTGAGTATG; reverse – GAAGAGCTACGGTTCGGTTAAG). 1 µl of T7EI (NEB) was then added to 19 µl of purified (QIAquick PCR Purification Kit, QIAGEN) and hybridised PCR product containing approximately 200 ng of the PCR product, digested for 15 minutes at 37°C, and immediately visualised on a 1% agarose electrophoresis stained with Midori Green Advance (Nippon). T7 endonuclease-mediated cleavage of mismatched base pairs at the target site is expected to yield products of approximately 518 and 226 bp.

### Molecular genotyping

To molecularly characterise HDR knock-in events, the genomic DNA (gDNA) of G_1_ and G_2_ mosquitoes fluorescently expressing the *actin5c::eCFP* construct was extracted as individual samples using the Wizard® Genomic DNA Purification Kit (Promega). Regions of gDNA were then amplified according to the three primer pairs as indicated in Fig. 1. Briefly, the primer pairs amplified three regions localised to the insertion site within the *white* gene: primer pair I (forward – GGTTAACGTATGCGGCAAACAC; reverse – GTGTCGCCACCATCTGTGGTAAG) flanked the full insertion region including the 5’ and 3’ homology arms, amplifying a product of 3985 bp in individuals containing the *white*^*CFP*^ allele or 2302 bp in wild-type individuals; primer pair II [forward – GGTTAACGTATGCGGCAAACAC (as in primer pair I); reverse – CGACAACCACTACCTGAGC], which spanned the 5’ region of the insertion and amplified a product of 1625 bp in *white*^*CFP*^ individuals, and primer pair III [forward – GCAGATGAACTTCAGGGTCAGC; reverse – GTGTCGCCACCATCTGTGGTAAG (as in primer pair II)] which spanned the 3’ region of the insertion and amplified a product of 1917 bp.

## ACKNOWLEDGEMENTS

This work was supported by pump-prime funding from ANTI-VeC (AV-PP20), a BBSRC/GCRF Network Grant. We are extremely grateful to Helen Williams and members of the Liverpool Insect Testing Establishment (LITE), in particular James Court and Amy Guy, and their expertise in rearing large populations of *An. funestus* colonies, which made this project a lot easier than it could have been. Matt Craske provided excellent technical assistance in the breeding of transgenic lines. We are also grateful to Leon Mugenzi, Andrew Hammond, Linta Grigoraki and Kyros Kyrou for helpful discussions on initial optimisation of injection technique and DNA cloning protocols.

## AUTHOR CONTRIBUTIONS

Conceptualization (TN, CW); Data Curation and Formal Analysis (CQ, AA, TN); Funding Acquisition (TM, CW); Investigation (TN, AA, CQ); DNA cloning (AA, TN); Transgenesis (CQ, TN); Molecular Genotyping (CQ); Methodology (AA, TN); Project Administration and Supervision (TN); Visualization (CQ, TN); Writing-original draft (CQ, TN); Writing – review and editing (CQ, AA, CW, TN).

## FIGURE LEGENDS

**Supplementary Figure 1 Detecting CRISPR-generated end-joining mutations in the soma using the T7 Endonuclease assay**. PCR amplicons (744 bp) from wild-type (wt_1_ and wt_2_) and representative G_0_ individuals exhibiting transient *white*^*CFP*^ expression from experiments A and C were digested with T7 endonuclease (+). The presence end-joining mutations at the CRISPR target site among the amplified fragments leads to mismatched base pairing that is susceptible to digestion by T7. Very partial digestion of the amplicon is evident in individuals A_m2_, A_m3_ and C_f1_, consistent with low overall levels of end-joining activity, while no digestion is evident in wild-type samples or individuals A_m1_ and C_f2_. None of the individuals tested showed eye colour mosaicism. Bands produced at approximately 518bp and 226bp are indicative of mismatches produced at or near the site of Cas9 cleavage. † - individual produced transgenic offspring.

