## Supplementary Figure 1 for "CRISPR-mediated knock-in of transgenes into the malaria vector *Anopheles funestus*"

### Slide 1
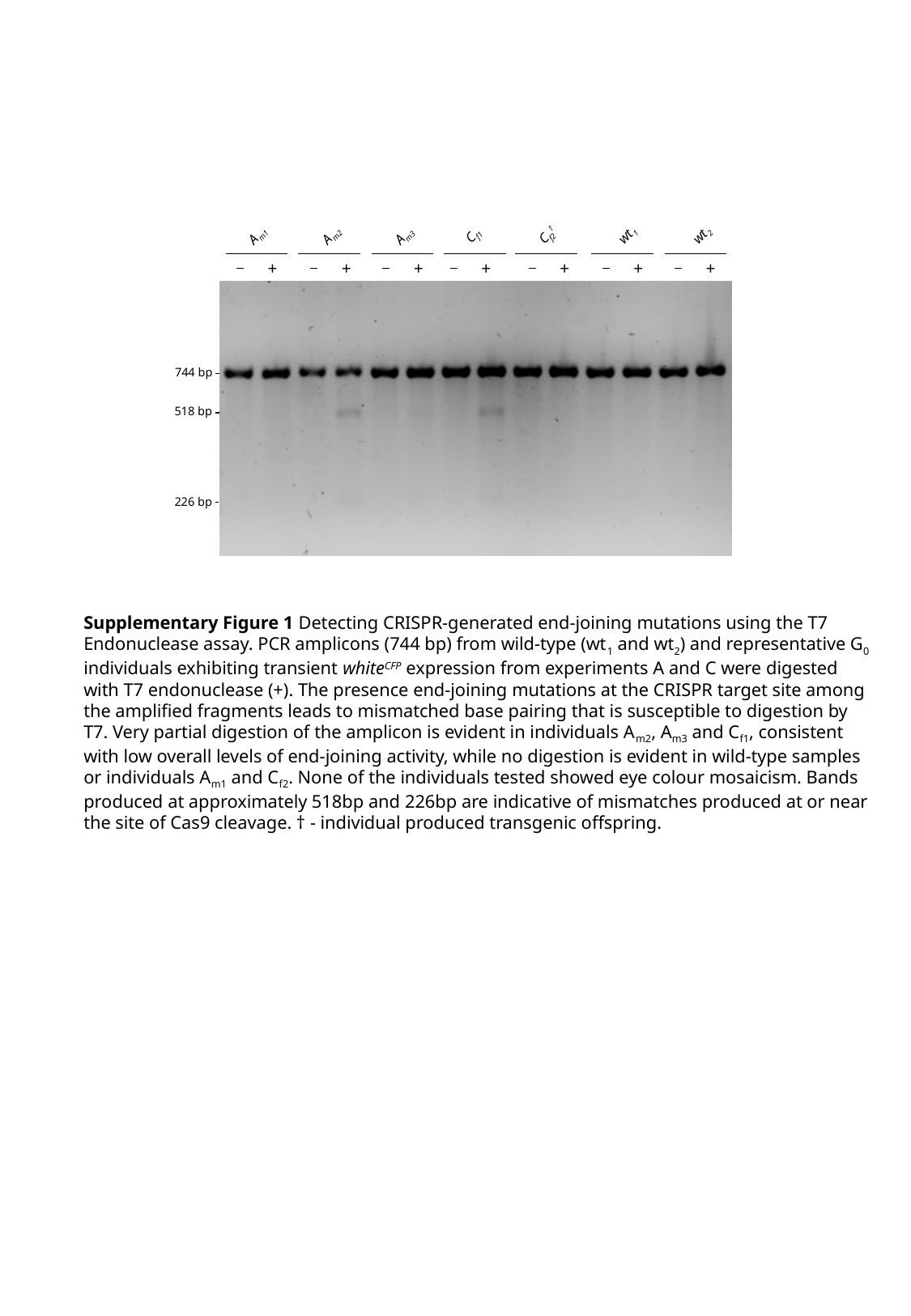

A_m2_
C_f2_^†^
C_f1_
A_m1_
H_2_O
wt_1_
wt_2_
A_m3_
+
+
+
+
+
+
+
+
744 bp
518 bp
226 bp
Supplementary Figure 1 Detecting CRISPR-generated end-joining mutations using the T7 Endonuclease assay. PCR amplicons (744 bp) from wild-type (wt_1_ and wt_2_) and representative G_0_ individuals exhibiting transient white^CFP^ expression from experiments A and C were digested with T7 endonuclease (+). The presence end-joining mutations at the CRISPR target site among the amplified fragments leads to mismatched base pairing that is susceptible to digestion by T7. Very partial digestion of the amplicon is evident in individuals A_m2_, A_m3_ and C_f1_, consistent with low overall levels of end-joining activity, while no digestion is evident in wild-type samples or individuals A_m1_ and C_f2_. None of the individuals tested showed eye colour mosaicism. Bands produced at approximately 518bp and 226bp are indicative of mismatches produced at or near the site of Cas9 cleavage. † - individual produced transgenic offspring.
